# Computational Design for Engineering Layered Tissue Architectures via Cell–Cell Interfacial Tension Modulation

**DOI:** 10.64898/2026.03.17.712503

**Authors:** Chayanit Thiticharoentam, Shuya Fukamachi, Shuhei A. Horiguchi, Satoru Okuda

**Affiliations:** Graduate School of Frontier Science Initiative, Kanazawa University, Kanazawa 920-1192, Japan; Nano Life Science Institute, Kanazawa University, Kanazawa 920-1192, Japan; Sapiens Life Sciences, Evolution and Medicine Research Center, Kanazawa 920-1192, Japan

**Keywords:** Tissue engineering, Cell sorting, Tissue morphogenesis, Stratification, 3D vertex model, Multicellular mechanics

## Abstract

The spatial arrangement of cells is fundamental to the mechanical and functional integrity of three-dimensional (3D) tissues, yet engineering spatially well-controlled tissue architectures remains challenging. Here, we computationally investigated how layered tissue architectures can be designed by modulating cell-cell interfacial tension. We performed simulations using a 3D vertex model and systematically varied interfacial tension magnitudes. The simulations generated a range of layered tissue architectures, including planar monolayers, bilayers, and structurally stratified states. In homogeneous cell populations, increasing interfacial tension drove transitions from monolayer to structurally stratified configurations. In heterogeneous populations, differential interfacial tensions induced out-of-plane cell sorting and the formation of compositionally sorted multilayers. Moreover, a recursive tension design strategy enabled hierarchical organization of multiple cell types into separate layers. Notably, this recursive scheme uses only two tension levels (high vs. low) assigned across interfaces and can, in principle, be extended to specify layered architectures with an arbitrary number of layers. Together, these results identify cell-cell interfacial tension as a tunable mechanical parameter for regulating layered tissue architecture and provide design principles for layered tissue engineering and regenerative medicine.

## 1. Introduction

Tissue engineering integrates biological principles with engineering strategies to fabricate artificial constructs that replicate the structure and function of natural tissues (Almeida et al., 2025; Cadavid et al., 2024; Peneda Pacheco et al., 2021; Zurina et al., 2020). Bottom-up approaches based on the assembly of living cells are increasingly explored as alternatives to scaffold-based methods for building three-dimensional (3D) tissue models with improved physiological relevance (Almeida-Pinto et al., 2023; Cadavid et al., 2024; Gaspar et al., 2020). However, functional outcomes are often limited by the difficulty of reproducing the multiscale architecture and mechanical behavior of complex tissues (Athanasiou et al., 2013; Gaspar et al., 2020; Kim et al., 2021; Zurina et al., 2020). Recent advances in cell-surface engineering provide tools to program cell-cell interactions and bias multicellular self-organization (Toda et al., 2019, 2018). Nevertheless, reliably controlling cellular positioning to achieve spatial organization and mechanical stability remains challenging, particularly for multilayered architectures (Almeida et al., 2025; Kim et al., 2021).

Biological tissues acquire diverse layered cellular architectures through intrinsic self-organization (Lee et al., 2022; Sasai, 2013; Sthijns et al., 2019). These architectures often emerge through structural stratification from an initially simple cell monolayer, but the resulting tissue structures can differ substantially in cellular composition and spatial arrangement. In addition, individual layers may themselves contain structurally stacked cells of similar identity. For instance, the skin maintains a stratified structure through coordinated cell turnover, in which basal progenitors continuously differentiate and migrate upward to replace differentiated cells at the surface (Yokouchi et al., 2016). During embryonic morphogenesis, such layered organizations are further refined by cytoskeletal reorganization, enabling cells to elongate and pack vertically into distinct tissue layers (Heller et al., 2014; Ipponjima et al., 2016). In contrast, the mammary gland establishes an ordered multilayered architecture in which cell sorting arranges distinct cell populations into inner and outer layers through differences in cortical contractility (Cerchiari et al., 2015). In parallel, synthetic approaches have demonstrated that programmed cell-cell interactions can drive multicellular self-organization into multilayered architectures in vitro (Toda et al., 2018). Nevertheless, reliably producing multilayered tissue architectures with prescribed cellular organization remains challenging.

The dynamic remodeling and spatial organization observed during tissue development are fundamentally shaped by the mechanical behaviors of individual cells, and often described in terms of effective cell surface tensions (Houtekamer et al., 2022; Lecuit and Lenne, 2007). At cell-cell contacts, intercellular surface tension emerges from a balance between intercellular adhesion, which promotes contact expansion, and cortical tension, which acts to minimize cell-cell interfaces (Heisenberg and Bellaïche, 2013; Kondo and Hayashi, 2015). Cortical tension is actively generated by contractile forces in the actomyosin cytoskeleton, whereas adhesion is mediated by adherens junctions that couple external contacts to the intracellular F-actin network (Burda et al., 2023; Campàs et al., 2024; Clarke and Martin, 2021; Heer and Martin, 2017; Mao and Baum, 2015). Spatial regulation of this mechanical interplay governs morphogenetic processes such as cell sorting, shape transitions, and geometric patterning, which are essential for the emergence of structured tissues (Cheng-Lin Lv and Bo Li, 2025; Fagotto, 2014; Lemke and Nelson, 2021). Despite the recognized importance of cell-cell interfacial tension, it remains unclear how modulation of this tension can be used as a design handle to program predictable layered tissue architectures.

Computational modeling enables systematic exploration of multicellular dynamics and provides a powerful framework for testing design principles of tissue architecture formation (Buttenschön and Edelstein-Keshet, 2020; Glen et al., 2019; Kim et al., 2020). A variety of frameworks, including agent-based, cellular Potts, phase-field, and Voronoi models, have been developed to simulate tissue morphogenesis across diverse contexts and levels of structural complexity (Brodland, 2004; Keshavanarayana et al., 2025; Lam et al., 2022; Sivakumar et al., 2022; Velazquez et al., 2018). For predicting layered tissue architecture formation while explicitly capturing cell deformability and direct cell-cell interactions, we employ a 3D vertex model, which enables simulation of structurally layered architectures and mechanically driven transitions under prescribed interfacial tensions (Briñas-Pascual et al., 2024; Fukamachi et al., 2024; Honda et al., 2004; Okuda et al., 2015a).

In this study, we investigate how layered tissue architectures can be designed by modulating interfacial tensions. Using a 3D vertex model, we perform numerical simulations to examine the emergence of layered structures and the underlying transition processes. In homotypic tissues composed of a single cell type, varying the magnitude of interfacial tensions yields either a monolayer or a structurally stratified state: low tensions yield a monolayer, whereas higher tensions drive a transition to structural stratification. In heterotypic tissues, we introduce a strategy for assigning differential tensions across interfaces, enabling control of cell-type spatial organization, including segregation into distinct layers or uniform intermixing depending on the relative tensions. Finally, we extend this approach with a recursive tension-design scheme for multi-cell-type systems, yielding compositionally sorted multilayers and hierarchical 3D architectures through controlled cell sorting. Together, these results establish interfacial tension as a tunable mechanical handle for designing layered tissue architectures.

## 2. Methods

### 2.1 3D vertex model framework

In the 3D vertex models, cells are assumed to be tightly packed such that the boundary between two adjacent cells is represented by a polygonal face (Honda et al., 2004; Honda and Nagai, 2015; Lange et al., 2025; Okuda et al., 2019, 2015a). Each cell is represented as a polyhedron whose geometry is defined by its boundary faces. Accordingly, the multicellular structure is described by a single network of the vertices and edges shared among neighboring cells.

The system box was defined as 0 ≤ *x* < *L*_*x*_, 0 ≤ *y* < *L*_*y*_, with the *z*-direction treated as unbounded (−∞ < *z* < ∞) in the *xyz*-coordinate system (Fig. 1a). Periodic boundary conditions were applied at *x* = 0, *L*_*x*_ and *y* = 0, *L*_*y*_. As an initial condition, *N*_T_ cells were arranged as a monolayer in the *xy*-plane within the box. The top and bottom free surfaces of the cell sheet along the *z*-direction were denoted as surfaces A and B, respectively (Fig. 1a and b). Faces shared by adjacent cells were referred to as lateral surfaces (Fig. 1b).

**Fig. 1.**
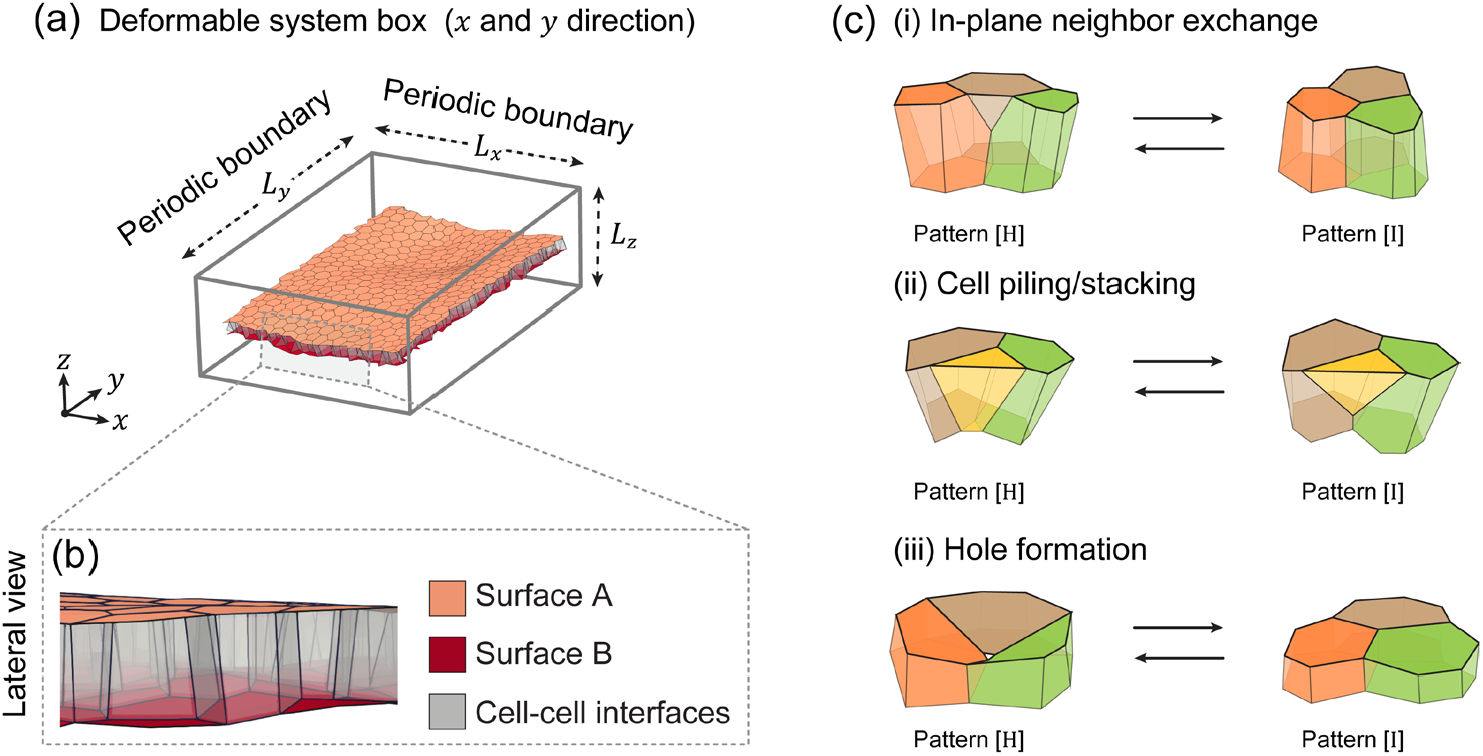
Modeling a monolayer cell sheet using the 3D vertex model. (a) Schematic of the monolayer cell sheet. Periodic boundary conditions are applied at *x* = 0, *L*_*x*_, *y* = 0 and *L*_*y*_. (b) Definition of tissue components: free surface A (orange), free surface B (red), and cell-cell interfaces (gray). (c) Examples of topological reconnection operations between [H] and [I] configurations used in the 3D vertex model: (i) in-plane neighbor exchange, (ii) cell piling/stacking, and (iii) hole formation.

Cell dynamics were calculated by tracking the motion of the network vertices. The time evolution of the *i* -th vertex location, denoted by ***r***_*i*_ (a 3D position vector of the *i* -th vertex), is given by the overdamped equation, following a previous study (Okuda et al., 2015b):

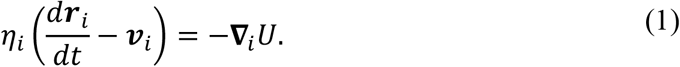

The left-hand side of Eq. (1) represents the viscous friction force exerted on the *i*-th vertex. Here, *η*_*i*_ is the friction coefficient, and ***v***_*i*_ is a local reference velocity around the *i*-th vertex. The right-hand side of Eq. (1) represents the mechanical force acting on the vertex, derived from the effective energy *U*. ∇_*i*_ denotes the gradient with respect to ***r***_*i*_.

The friction coefficient *η*_*i*_ was assumed to arise from frictional interactions between adjacent cells and was modeled as

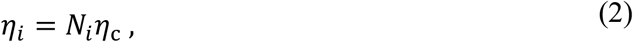

where *N*_*i*_ is the number of cells adjacent to the *i*-th vertex, and constant *η*_c_ is the intercellular friction coefficient.

The local reference velocity ***v***_*i*_ was defined as the average of the velocities of the cells adjacent to the *i*-th vertex:

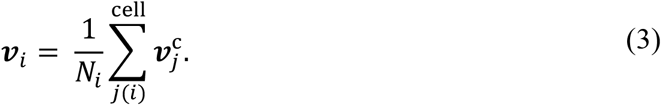

where the summation denotes the set of cells sharing the *i*-th vertex. The velocity of the *j*-th cell 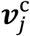 was calculated as the average velocity of the vertices constituting the *j*-th cell.

During cell motion, individual edges and polygonal faces in the network occasionally shrink. When the length of a relevant edge fell below a prescribed threshold, the topological network was reconnected via [I]-to-[H] and [H]-to-[I] operations (Honda et al., 2004) (Fig. 1c). These operations implement 3D topological transitions that represent cellular rearrangements, cell stacking, and hole formation, according to predefined rules (Okuda et al., 2013; Okuda and Fujimoto, 2020).

### 2.2 Effective energy function

The effective energy of the cell sheet is defined as

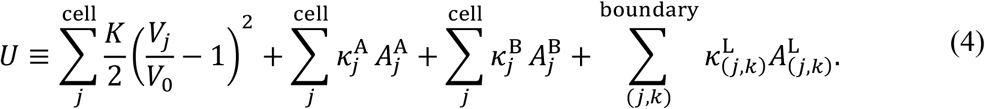

The first term represents the volume elastic energy, where *K* is the elastic modulus (chosen large to enforce near-incompressibility), *V*_*j*_ is the volume of the *j*-th cell, and *V*_0_ is the reference volume. The second and third terms represent the surface energies associated with top (surface A) and bottom (surface B) free surfaces of the sheet: 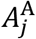 and 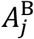 are the areas of the corresponding surfaces of the *j*-th cell, and 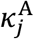 and 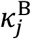 are the associated surface tensions. The final term represents the energy of the cell-cell interface, where 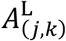 is the area of the interface shared by the adjacent *j*-th and *k*-th cells, and 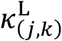 is the corresponding cell-cell interfacial (lateral) tension.

### 2.3 Stress-free periodic boundary condition

The axial stress in the *xy*-plane was assumed to be approximately free; the periodic boundary was allowed to deform affinely by dynamically adjusting the box lengths *L*_*x*_ and *L*_*y*_. The time evolution of *L*_*α*_ (*α* ∈ {*x, y*}) was described by

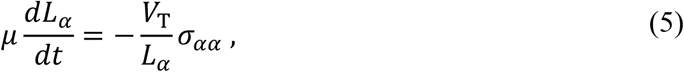

where constant *μ* is the effective friction coefficient of the tissue with respect to its surroundings, and *σ*_*αα*_ is the normal component of the stress tensor of the system along the *α*-th axis. Constant *V*_T_ is the total cell volume, given by

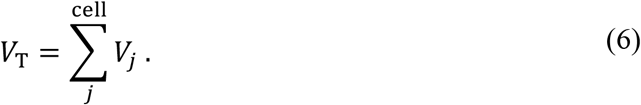

The stress tensor, denoted by *σ*_*αα*_, is given by

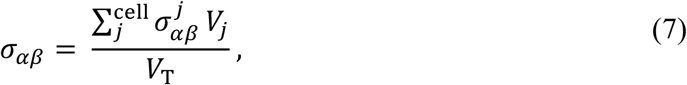

where 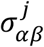 denotes the stress tensor of the *j*-th cell. The cell stress tensor 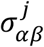 was defined by

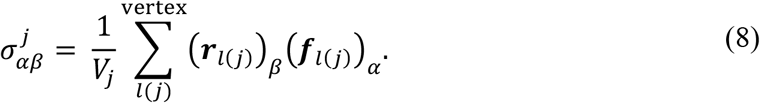

where the summation is taken over all vertices of the *j* -th cell. Here, ()_*α*_ indicates the *α* -th component of a vector. The vector ***r***_*l*(*j*)_ is the distance vector from the center of the *j*-th cell to the position of the *l*-th vertex, and ***f***_*l*(*j*)_ is the force acting on that vertex. The vertex force ***f***_*l*(*j*)_ was obtained from the effective energy of the *j*-th cell, *u*_*j*_, as ***f***_*l*(*j*)_ = −*du*_*j*_/*d****r***_*l*_. Here, *u*_*j*_ is defined as

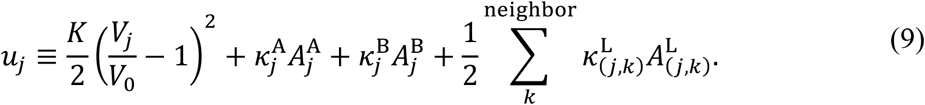

The factor 1/2 avoids double counting of the cell-cell interfacial contributions shared by neighboring cells.

### 2.4 Parameter setting for surface tension in two-cell-type

In the two-cell-type simulations, each cell was assigned a type label *c*_*j*_ ∈ {I, II}. The free-surface tension coefficients in the effective energy (Eq. (4)), 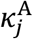 and 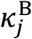, were allowed to depend on cell type. In principle, a heterogeneous tissue admits many independent tension parameters (type-dependent tensions on each free surface and type-pair-dependent cell-cell interfacial tensions), which makes systematic exploration impractical. We therefore restricted the free-surface tensions to two parameters, *κ*_1_ and *κ*_2_, while fixing all lateral interfacial tensions to *κ*_0_ as a reference.

Specifically, we imposed the free-surface tensions as

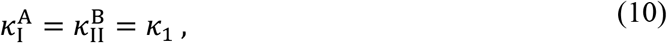

and

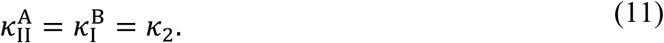

where 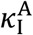 denotes the free-surface tension coefficient on surface A for cells of type I, and 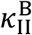 denotes the free-surface tension coefficient on surface B for cells of type II (and similarly for the other combinations). Thus, *κ*_1_ and *κ*_2_ are the two control parameters that prescribe how the two cell types preferentially interact with the two free surfaces.

For cell–cell interfaces, we fixed the lateral interfacial tension to a single baseline value:

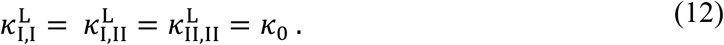

Here, 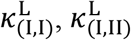, and 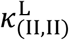 denote the cell-cell (lateral) interfacial tension coefficients between two type-I cells, a type-I and a type-II cell, and two type-II cells, respectively.

### 2.5 Parameter settings for recursive tension design in multi-cell-type systems

To establish a tractable strategy for designing interfacial tensions to engineer multilayered structures in multi-cell-type systems, we proposed a recursive approach. Specifically, the tension coefficients in the effective energy (Eq.(4))—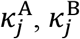, and 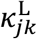—were assigned either *κ*_3_ or *κ*_4_, according to the cell-type combinations. The scheme was systematically applied to systems containing *M* distinct cell types (*M* = 2,3,4,5). Let the cell type of the *j*-th cell be denoted by *c*_*j*_ ∈ 𝒞, where 𝒞 = I, II, …, *M* is the ordered set of cell types. For mathematical formulation, we introduce an ordinal function *n*(*c*) that maps each Roman numeral to its corresponding integer value (e.g., *n*(I) = 1, *n*(II) = 2, …, *n*(*M*) = *M*).

#### Free surface tensions

To favor the positioning of specific cell types at the free surfaces, we assigned *κ*_4_ to the target cell type and *κ*_3_ to all other types. The surface-A tension of the *j*-th cell was defined as:

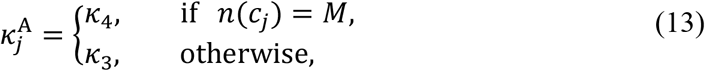

and the surface-B tension was defined as:

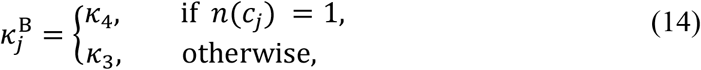

When *κ*_3_ > *κ*_4_, these assignments minimize the energy when type *M* is exposed to surface A and type I is exposed to surface B, thereby biasing their localization toward the corresponding free surfaces.

#### Cell-cell interfacial (lateral) tension

To model cell sorting, we defined the interfacial tension, 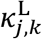 between the neighboring *j*-th and *k*-th cells. This interaction depends on the compatibility of their respective cell types, *c*_*j*_ and *c*_*k*_:

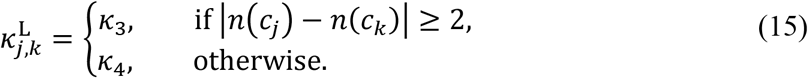

The choice of whether to denote these tensions by *κ*_3_ and *κ*_4_ is arbitrary; what matters is how they are assigned to the two interface classes. Under this formulation, interfaces between identical types *c*_*j*_ = *c*_*k*_ or immediate ordinal neighbors (e.g., type I and II) are assigned to *κ*_4_. Conversely, interfaces between non-adjacent types (e.g., type I and III) are assigned to *κ*_3_. This assignment penalizes contact between non-adjacent steps in the hierarchy relative to the reference state, driving the system toward a sorted configuration.

### 2.6 Initial conditions and nondimensional values

The initial conditions were prepared as follows:

#### Single cell type

A monolayer cell sheet of 401 cells of a single type. Cells were aligned on the *xy*-plane.

#### Two cell types

A monolayer cell sheet of 600 cells of the two cell types (I and II). Cells were randomly distributed to achieve an approximately equal ratio (1:1).

#### Three cell types

A monolayer cell sheet of 900 cells of the three cell types (I, II, and III) was randomly distributed to achieve an approximately equal ratio (1:1:1).

#### Four cell types

A monolayer cell sheet of 1,200 cells of the four cell types (I, II, III, and IV) was randomly distributed to achieve an approximately equal ratio (1:1:1:1).

#### Five cell types

A monolayer cell sheet of 1,500 cells of the five cell types (I, II, III, IV, and V) was randomly distributed to achieve an approximately equal ratio (1:1:1:1:1).

To solve Eq. (1), we applied two distinct nondimensionalization schemes depending on the system.

#### For single- and two-cell-type systems

Parameters were nondimensionalized using the unit length *l* = (*V*_0_)^1/3^, the unit energy *e* = 5*κ*_0_(*V*_0_)^2/3^ and the unit time *τ* = 0.8*η*_c_/*κ*_0_. The normalized values of the corresponding parameters were set to *V*_0_ = *l*^3^, *κ*_0_ = 0.2*e*/*l*^2^, and *η*_c_ = 0.25*τe*/*l*^2^.

#### For recursive tension design systems

Parameters were nondimensionalized using the unit length *l* = (*V*_0_)^1/3^, the unit energy *e* = 10*κ*_4_(*V*_0_)^2/3^ and the unit time *τ* = 0.4*η*_c_/*κ*_4_. The normalized values of the corresponding parameters were set as *V*_0_ = *l*^3^, *κ*_4_ = 0.1*e*/*l*^2^, and *η*_c_ = 0.25*τe*/*l*^2^.

### 2.7 Numerical procedures and statistics

The numerical integration of Eq. (1) was carried out using Euler’s method with a time step of Δ*d* (= 0.005 *τ*). At each time step, the vertex velocities were determined implicitly through an iterative convergence procedure until the mean residual error fell below *RE*_th_ (= 0.001 *e*/*l*). During this process, topological operations were performed on edges and triangular faces whenever their lengths fell below Δ*l*_th_ (= 0.05 *l*), at every time interval Δ*d*_r_ (= 1.0 *τ*). The numerical parameters used in these simulations are summarized in Table 2. This procedure ensured that the system reached an energy-minimized configuration for each parameter set.

For each parameter set, three independent simulations were performed using different initial monolayer cell sheets generated with distinct random seeds. Means and standard deviations of the observed quantities were calculated from these three simulations.

### 2.8 Evaluations of tissue structure

The tissue structure was evaluated using the following two quantities.

#### Average stratification number (N_l_)

The average number of cells stacked along the tissue thickness, *N*_l_, was defined as

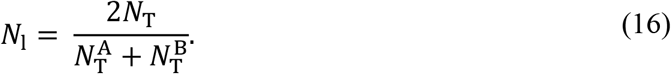

This definition corresponds to the trapezoidal formula used to measure the average tissue height (Martínez-Ara et al., 2022). Here, *N*_T_ denotes the total number of cells, while 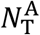 and 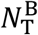 denote the total number of cells that possess free tissue surfaces of type A and type B, respectively.

#### *Tissue thickness* (*T*)

Tissue thickness, represented by *T*, was defined as

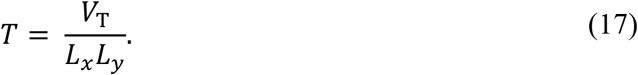

where *V*_T_ is the total volume of cells, and *L*_*x*_ and *L*_*y*_ denote the lengths of the system box in the *x*- and *y*-directions, respectively.

## 3. Results

### 3.1 Mechanical control of single-cell-type tissue architecture through free-surface tension

To explore how free-surface tension can be leveraged to control tissue architecture in a homogeneous tissue, we simulated a cell sheet composed of a single cell type (Fig. 2a, Sec. 2.6). As shown in Fig. 2a-(i), the free-surface tension coefficients on surfaces A and B were denoted by *κ*^A^ and *κ*^B^, respectively. The cell-cell interfacial (lateral) tension between neighboring cells, *κ*_0_ (Fig. 2a-(ii)), was used as the reference tension scale. Because exchanging surfaces A and B yields an equivalent system, the results for *κ*^A^ and *κ*^B^ can be mapped onto each other by swapping the labels A and B. Therefore, simulations were performed only for *κ*^A^ ≤ *κ*^B^, and the remaining region (*κ*^A^ > *κ*^B^) was obtained from the corresponding results with A and B exchanged.

**Fig. 2.**
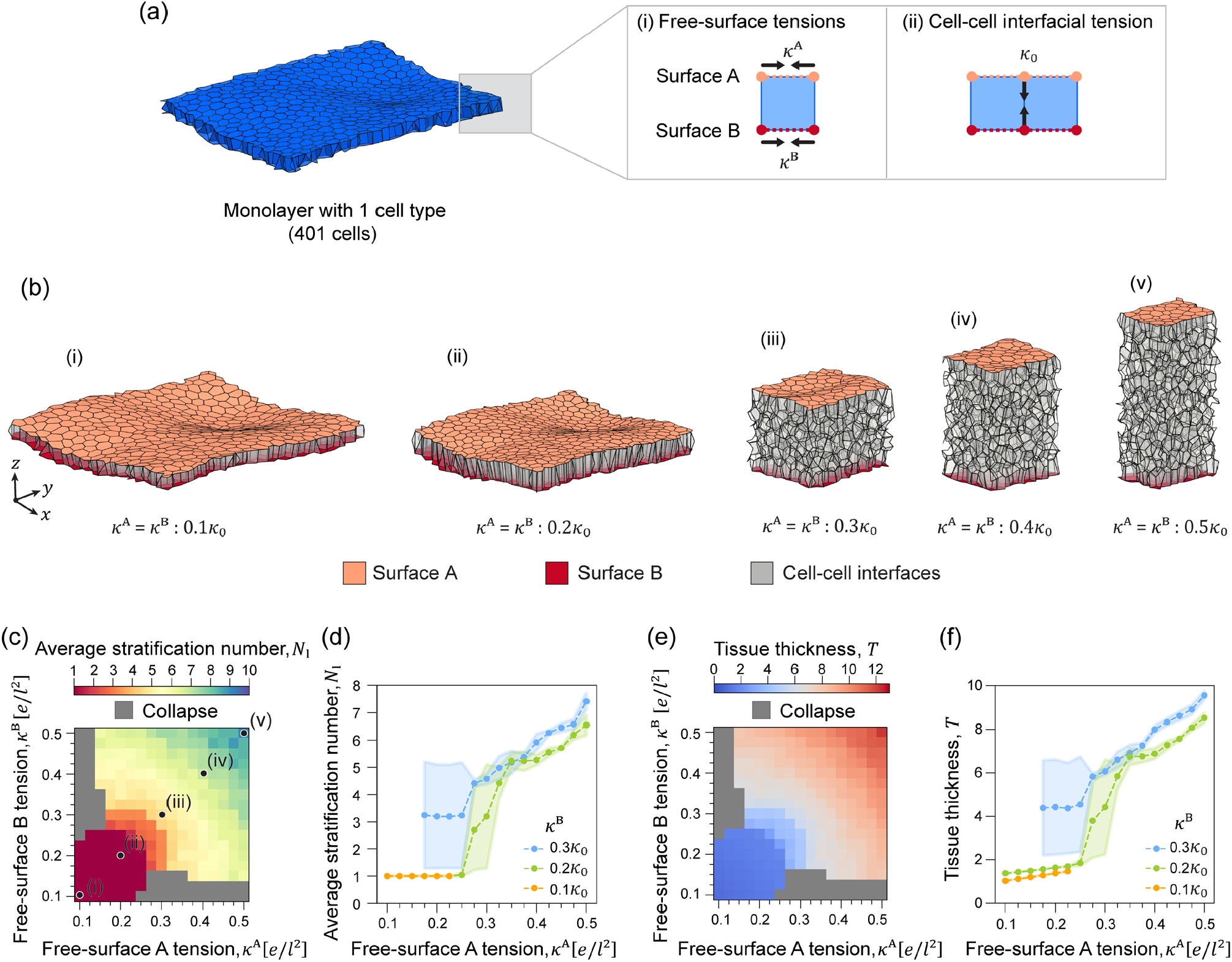
Single-cell-type tissue architecture as a function of free-surface tensions. (a) Initial condition and parameter definitions for a monolayer sheet of cells (blue). (i) The free-surface tension coefficients in the effective energy (Eq. (4)) were set to 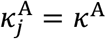 and 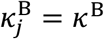. (ii) The lateral cell-cell interfacial tension is set to 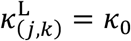. (b) Snapshots under the condition of equal free-surface tensions, *κ*^A^ = *κ*^B^: (i) 0.1*κ*_0_, (ii) 0.2*κ*_0_, (iii) 0.3*κ*_0_, (iv) 0.4*κ*_0_, and (v) 0.5*κ*_0_, shown at *d* = 2000*τ*. (c,d) Average stratification number *N*_l_ as a function of *κ*^A^ and *κ*^B^. (e,f) Average tissue thickness *T* as a function of *κ*^A^ and *κ*^B^. In (d,f), orange/green/blue curves indicate *κ*^B^ = 0.1*κ*_0_, 0.2*κ*_0_, and 0.3*κ*_0_, respectively. Simulations were performed only for the region for *κ*^A^ ≤ *κ*^B^; the results for *κ*^A^ > *κ*^B^ were obtained by exchanging *κ*^A^ and *κ*^B^.

We first examined the special case of equal free-surface tensions, *κ*^A^ = *κ*^B^, which is specifically highlighted in Fig. 2b. Under this condition, increasing *κ*^A^ (= *κ*^B^) led to a progressive increase in tissue thickness within a monolayer-like state (Fig. 2b-(i-ii)), followed by a transition to a structurally stratified state with smooth free surfaces (Fig. 2b-(iii-v)).

To determine design conditions for monolayer versus structurally stratified architectures, we next explored the (*κ*^A^, *κ*^B^) space by varying *κ*^A^ and *κ*^B^ independently and quantified the average stratification number *N*_l_ and the average tissue thickness *T* (Fig. 2c-f). When both *κ*^A^ and *κ*^B^ were small (approximately *κ*^A^ ≤ 0.2*κ*_0_ and *κ*^B^ ≤ 0.2*κ*_0_), the tissue remained in a monolayer state (*N*_l_ ≈ 1; Fig. 2c), while the monolayer thickness increased as *κ*^A^ and *κ*^B^ increased within this regime (Fig. 2e). In contrast, when the free-surface tensions exceeded this range, the system underwent a transition to a structurally stratified architecture (Fig. 2c), accompanied by further thickening (Fig. 2e). These results indicate that tuning the free-surface tensions provides a direct mechanical handle to switch homogeneous tissues between monolayer and structurally stratified organizations.

The observed transition can be interpreted from the competition between energetic contributions in Eq. (4). Increasing *κ*^A^ and *κ*^B^ increases the energetic cost of maintaining large free-surface areas, favoring configurations that reduce free-surface area for a given tissue volume. Structural stratification provides one such route by redistributing volume in the out-of-plane direction, although it can increase lateral interfacial area and associated cell-cell interfacial energy. The transition therefore emerges when the free-surface energy penalty becomes sufficiently large relative to the cell-cell interfacial contribution characterized by *κ*_0_, consistent with the threshold-like boundary observed in the (*κ*^A^, *κ*^B^) map (Fig. 2c).

These simulations show that free-surface tensions *κ*^A^ and *κ*^B^, relative to the cell-cell interfacial tension scale *κ*_0_, provide a practical control handle for switching a homogeneous cell sheet between monolayer and structurally stratified organizations. In the (*κ*^A^, *κ*^B^) design map, a monolayer regime persists for sufficiently small free-surface tensions, whereas larger values lead to a transition to structurally stratified architectures accompanied by tissue thickening. Mechanistically, this transition is attributed to a mechanical instability of the cell monolayer (Fukamachi et al., 2024; Okuda and Fujimoto, 2020); here, we focus not on the mechanism itself but on leveraging this instability as a design principle for engineering tissue architecture. More broadly, this design-oriented interpretation is consistent with prior experimental and modeling studies in which modulation of tension has been implicated in driving epithelial shape changes and tissue morphogenesis (Rozman et al., 2020; Sui et al., 2018).

### 3.2 Mechanical control of two-cell-type tissue architecture through free-surface tensions

To extend the design framework to heterogeneous tissues, we simulated a monolayer sheet composed of two cell types (types I and II) with a total of 600 cells (Sec. 2.6; Fig. 3a). The free-surface tension coefficients on surfaces A and B were parameterized using two values, *κ*_1_ and *κ*_2_, and assigned to each cell type according to Eqs. (10) and (11), i.e., 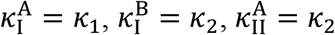, and 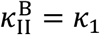 (Fig. 3a-(i)). The lateral (cell-cell interfacial) tension between neighboring cells was set to *κ*_0_ and used as the unit of tension (Fig. 3a-(ii); Table 1). We then explored the (*κ*_1_, *κ*_2_) parameter space to determine how engineered free-surface tensions control tissue architecture and cell-type organization. Because exchanging the labels of the two cell types maps (*κ*_1_, *κ*_2_) to (*κ*_2_, *κ*_1_), simulations were performed only for *κ*_1_ ≤ *κ*_2_, and the results for *κ*_1_ > *κ*_2_ were obtained by exchanging the cell-type labels.

**Table 1.**
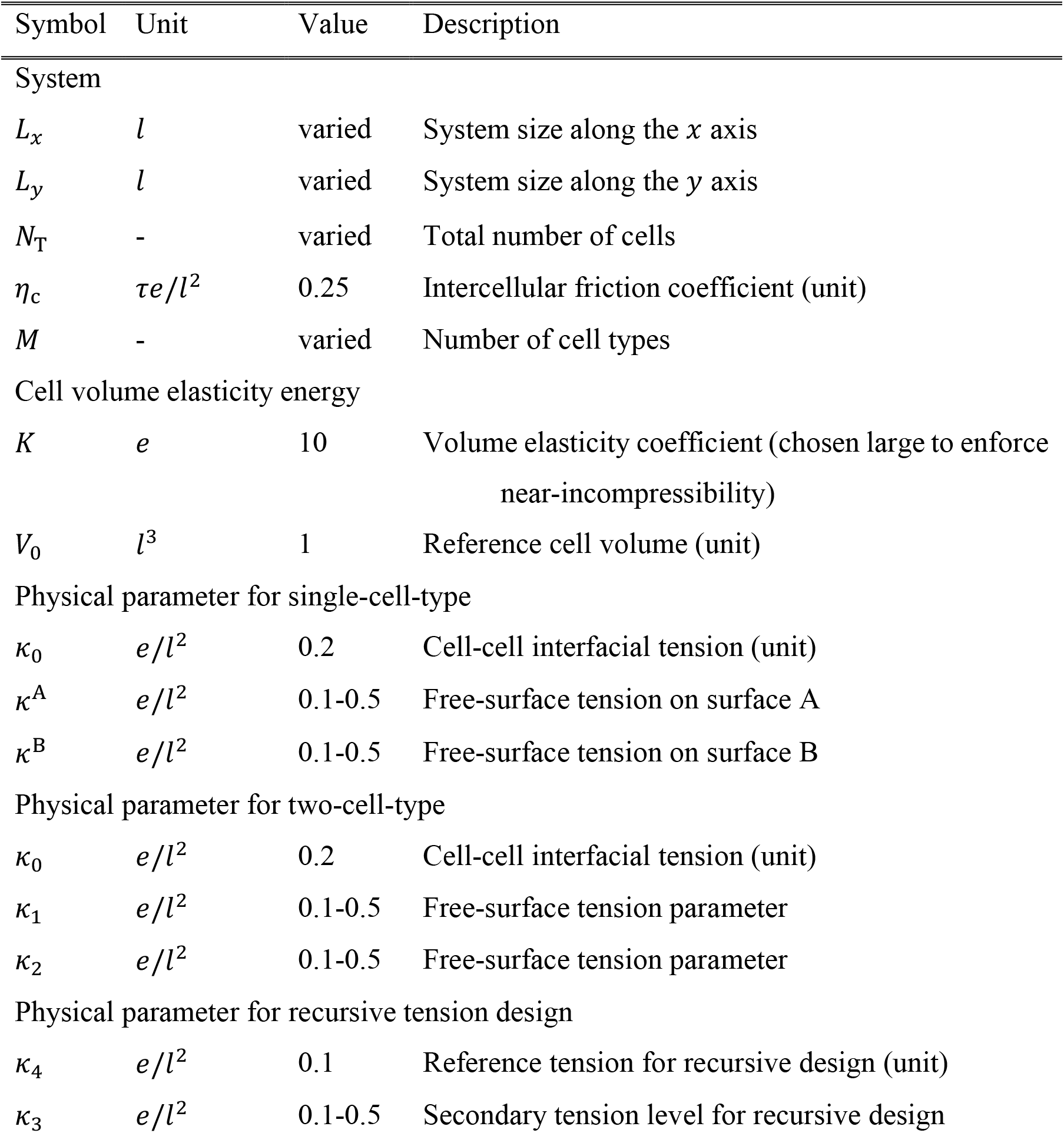
Physical parameters.

**Table 2.** Numerical parameters used in the model.

| Symbol | Unit | Value | Description |
| --- | --- | --- | --- |
| $\Delta t$ | $\tau$ | 0.005 | Time step for integrating vertex dynamics |
| $\Delta t_r$ | $\tau$ | 1.0 | Time interval for network reconnection updates |
| $\Delta l_{\text{th}}$ | $l$ | 0.05 | Edge-length threshold for triggering reconnections |
| $RE_{\text{th}}$ | $e/l$ | 0.001 | Threshold for the residual (convergence criterion) |

**Fig. 3.**
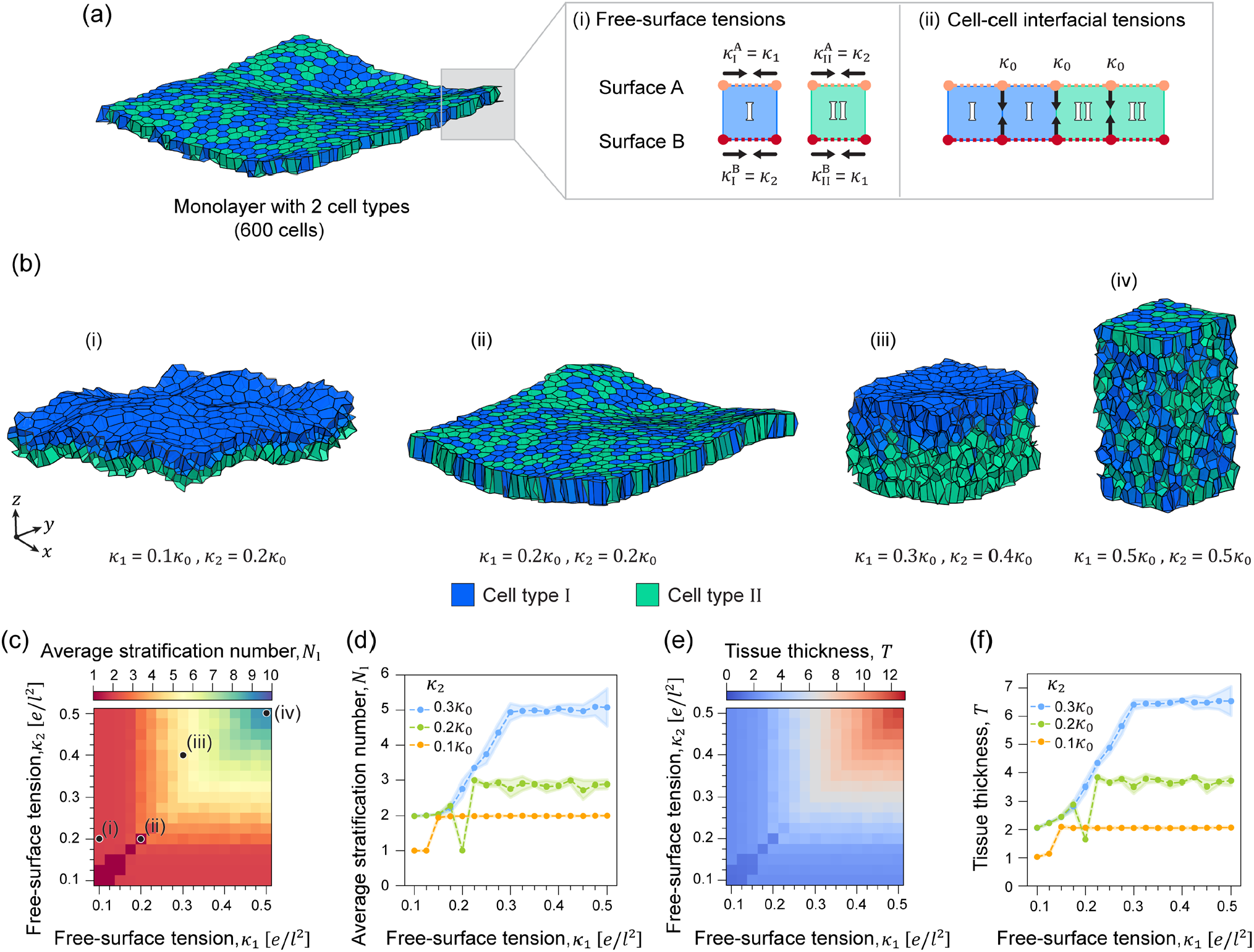
Two-cell-type tissue architecture as a function of engineered free-surface tensions. (a) Initial condition and parameter definitions for a monolayer sheet containing cell types I (blue) and II (green), initially mixed at random. (i) Free-surface tensions were assigned according to Eqs. (10) and (11): 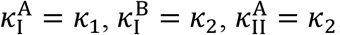 and 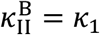. (ii) The lateral interfacial tensions were fixed according to Eq. (12): 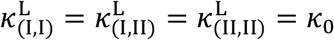. (b) Snapshots at *d* = 2000*τ* for (i) *κ*_1_ = 0.1*κ*_0_, *κ*_2_ = 0.2*κ*_0_; (ii) *κ*_1_ = 0.2*κ*_0_, *κ*_2_ = 0.2*κ*_0_; (iii) *κ*_1_ = 0.3*κ*_0_, *κ*_2_ = 0.4*κ*_0_; and (iv) *κ*_1_ = 0.5*κ*_0_, *κ*_2_ = 0.5*κ*_0_ . (c,d) Average stratification number *N*_l_ as a function of *κ*_1_ and *κ*_2_. (e,f) Average tissue thickness *T* as a function of *κ*_1_ and *κ*_2_ . In (d,f), orange/green/blue curves indicate *κ*_2_ = 0.1*κ*_0_, 0.2*κ*_0_, and 0.3*κ*_0_, respectively.

We first summarize representative tissue morphologies obtained by scanning (*κ*_1_, *κ*_2_) (Fig. 3b). When *κ*_1_ ≠ *κ*_2_, the two cell types undergo vertical sorting, producing a compositionally sorted multilayer in which one type preferentially occupies the top side and the other occupies the bottom side (sorted regime; Fig. 3b-(i,iii)). In contrast, when *κ*_1_ = *κ*_2_, the tissue remains largely unsorted and forms an architecture similar to the single-cell-type case under the same overall tension level (unsorted regime; Fig. 3b-(ii,iv)). Increasing both *κ*_1_ and *κ*_2_ thickens the tissue and can induce a transition from a thin sheet to a structurally stratified architecture, while the difference ∣ *κ*_1_ − *κ*_2_ ∣ primarily determines whether the resulting structure forms a compositionally sorted multilayer or remains compositionally mixed.

To quantify these trends, we measured the average stratification number *N*_l_ and the average tissue thickness *T* over independent simulations with different initial cell arrangements (Fig. 3c-f). In the low-tension region (approximately *κ*_1_ ≤ 0.2*κ*_0_ and *κ*_2_ ≤ 0.2*κ*_0_), the tissue adopted a bilayer organization with *N*_l_ close to 2 and a thickness that remained nearly constant (Fig. 3c and e; Eqs. (16) and (17)). Along one-dimensional slices of the map (Fig. 3d and f), the behavior depends on *κ*_2_: for small *κ*_2_, the tissue saturates in a bilayer state even as *κ*_1_ increases, whereas for larger *κ*_2_ the tissue exhibits a transition to compositionally sorted multilayers with progressively increasing structural stratification within individual layers once *κ*_1_ exceeds a condition-dependent boundary. These results demonstrate that tuning *κ*_1_and *κ*_2_ provides a practical mechanical handle to control the transition from a bilayer state to compositionally sorted multilayers, in which individual layers become increasingly structurally stratified, in two-cell-type tissues, with only small variations across independent initial conditions (Fig. 3d and f).

Finally, to characterize how cell types are distributed within multilayered tissues, we analyzed the composition of individual layers for representative multilayer states (Fig. 4). In a three-layer configuration (Fig. 4a), the topmost layer consisted almost exclusively of cell type I, the bottommost layer consisted almost exclusively of cell type II, and the middle layer contained a mixture of both types (Fig. 4b and c). In a five-layer configuration (Fig. 4d), the topmost and bottommost layers again became nearly pure layers of types I and II, respectively, whereas the interior layers exhibited mixed compositions with a clear gradient: the second layer from the top was enriched in type I, the middle layer was closer to an even mixture, and the second layer from the bottom was enriched in type II (Fig. 4e and f). Together, these analyses show that engineered free-surface tensions control not only overall tissue thickness and layering, but also the internal segregation pattern of cell types within multilayered architectures.

**Fig. 4.**
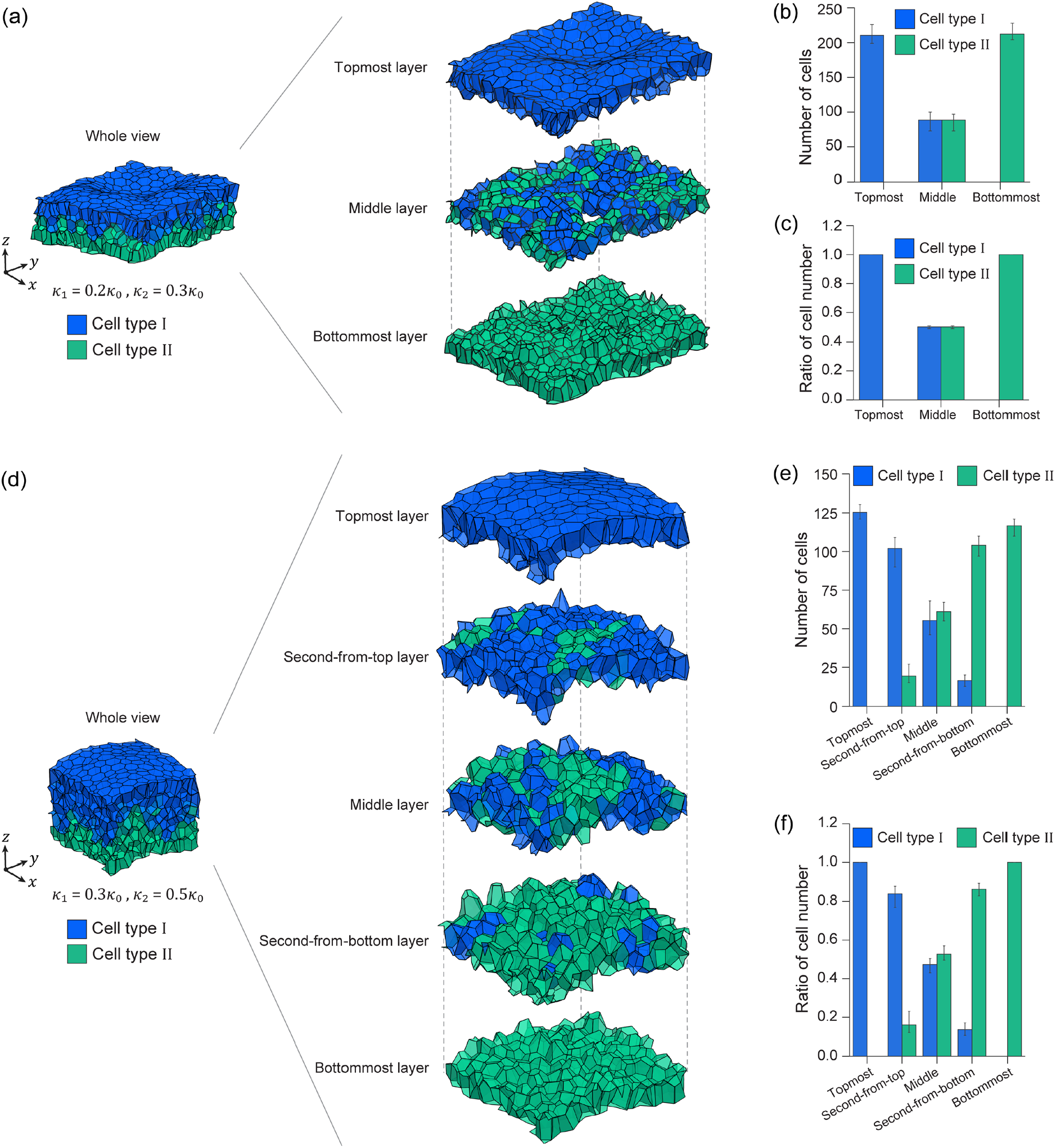
Layer composition and vertical segregation in two-cell-type multilayers. (a) Example of a three-layer structure formed by cell types I (blue) and II (green), with layers labeled as topmost, middle, and bottommost. (b) Cell counts in each layer for both cell types. (c) Fractional composition of each layer (cell count of each type divided by the total cell count in the layer). (d) Example of a five-layer structure, with layers labeled as topmost, second-from-top, middle, second-from-bottom, and bottommost. (e) Cell counts in each layer for both cell types. (f) Fractional composition of each layer. Snapshots in (a) were obtained at *κ*_1_ = 0.2*κ*_0_, *κ*_2_ = 0.3*κ*_0_ and *d* = 2000*τ*. Snapshots in (d) were obtained at *κ*_1_ = 0.3*κ*_0_, *κ*_2_ = 0.5*κ*_0_ and *d* = 2000*τ*. Error bars indicate standard errors.

These simulations support the view that strategic tuning of *κ*_1_ and *κ*_2_ can control both vertical sorting and multilayer formation in heterogeneous tissues. The emergence of sorted configurations when *κ*_1_ ≠ *κ*_2_ is consistent with the differential interfacial tension hypothesis, in which tissues minimize effective interfacial energies to segregate cell populations (Keister et al., 2024). Moreover, beyond binary sorting, our tension design produces well-defined tissue-scale interfaces and compositional gradients in multilayers, aligning with geometric and compositional features reported in other 3D tissue modeling studies (Sahu et al., 2021).

### 3.3 Mechanical control of multi-cell-type tissue architecture through recursive tension design

Building on our results for single- and two-cell-type tissues (Secs. 3.1-3.2), we next tested whether compositionally sorted multilayers can be generated in systems containing more than two cell types by prescribing tension coefficients in a rule-based manner. Specifically, we applied the recursive assignment scheme described in Sec. 2.5, in which the free-surface tensions 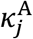 and 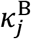 and the cell– cell boundary tensions 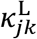 in Eq. (4) take one of two values, *κ*_3_ or *κ*_4_, depending on cell type and interface identity (Fig. 5a).

**Fig. 5.**
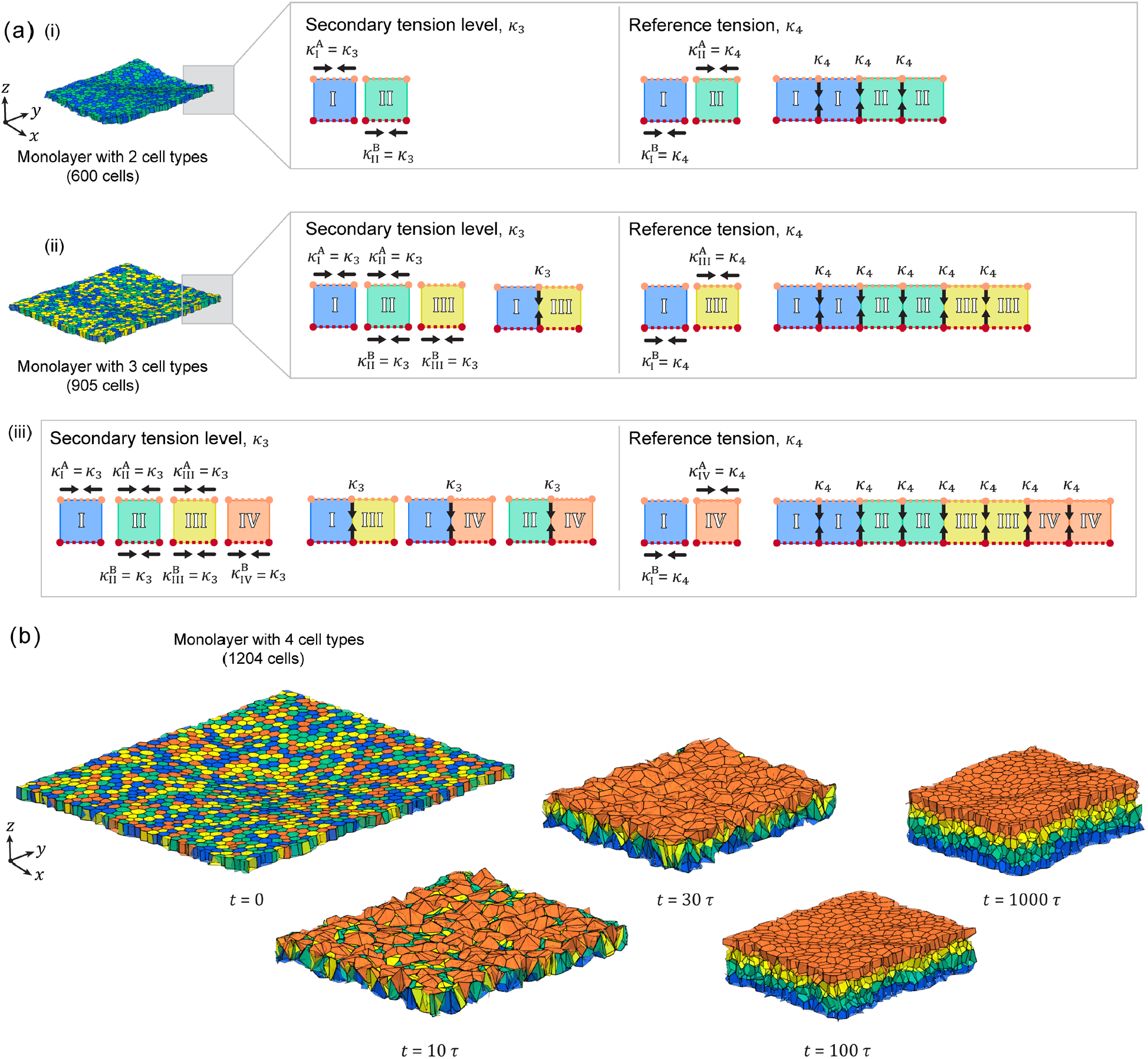
Recursive tension design and initial conditions for multi-cell-type systems. (a) Initial conditions and parameter assignments for multi-cell-type systems. Free-surface tensions 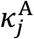 and 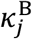 and cell-cell interfacial tensions 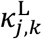 in the effective energy (Eq. (4)) were assigned as either *κ*_3_ or *κ*_4_ according to the recursive rules (Eqs. (13)-(15); Sec. 2.5). (b) Time evolution of the four-cell-type system showing the transition from an initially mixed monolayer to a stratified, spatially organized architecture under the representative condition *κ*_3_ = 0.4*κ*_4_. Cell types are indicated by distinct colors.

Simulations were performed for randomly mixed initial conditions containing *M* = 2 to 5 cell types (Sec. 2.6; Fig. 5a). We fixed *κ*_4_ as the reference tension and varied *κ*_3_ over the range 0.1*κ*_4_ ≤ *κ*_3_ ≤ 0.5*κ*_4_ (Sec. 2.7; Table 1). For each *M*, the value of each tension component was assigned to either *κ*_3_ or *κ*_4_ according to the recursive rules (Sec. 2.5), yielding a consistent two-level tension specification across different numbers of cell types (Fig. 5a).

To validate the recursive tension design proposed for multi-cell-type systems, we simulated the spatiotemporal evolution of a four-cell-type system (Fig. 5b). The simulation results illustrate the spontaneous transition from a randomized planar configuration to a compositionally sorted multilayer with structural stratification within individual layers. Initiated from a random, monolayer distribution at *d* = 0, the assembly undergoes significant rearrangement and out-of-plane remodeling by *d* = 30*τ*. As the simulation progresses (*d* = 100*τ* to *d* = 1000*τ*), distinct vertical segregation occurs, in which cell populations sort into discrete layers with cell types I through IV arranged sequentially from the bottommost to the topmost layer.

The morphology map for multi-cell-type systems shows that at the lowest value *κ*_3_ = 0.1*κ*_4_, the tissue remained close to a monolayer-like structure with an unsorted pattern (Fig. 6a). As *κ*_3_ increased, the initially mixed cell populations reorganized into spatially ordered configurations in which cell types formed distinct domains and compositionally sorted multilayers (Fig. 6a). The resulting spatial ordering of cell types depended on *M* and emerged from the interfacial-tension pattern prescribed by the recursive design rule. For example, in the three-cell-type system, type II occupied an internal region while types I and III localized toward opposite sides. In the four- and five-cell-type systems, additional types formed further separated domains consistent with the intended hierarchy, yielding progressively more complex compositionally sorted multilayers (Fig. 6a).

**Fig. 6.**
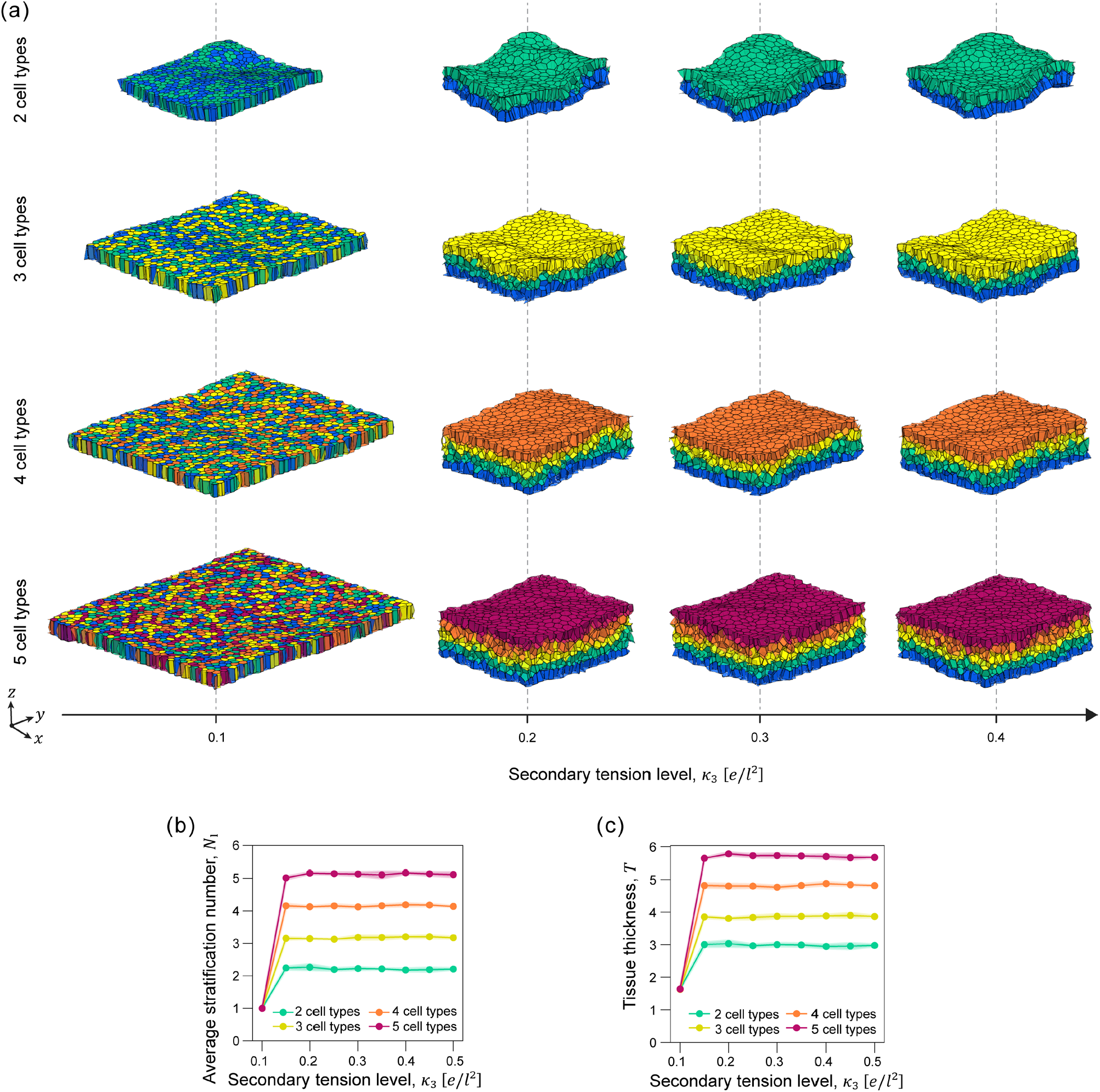
Multi-cell-type tissue morphologies as a function of secondary tension level for recursive design. (a) Morphology map for systems with two to five cell types (top to bottom). All snapshots are shown at *d* = 1000*τ* with *κ*_4_ = 0.1. (b) Average stratification number *N*_l_ as a function of *κ*_3_. (c) Average tissue thickness *T* as a function of *κ*_3_. Curves correspond to two (green), three (yellow), four (orange), and five (red) cell types.

To quantify these structural changes, we measured the average stratification number *N*_l_ and the average tissue thickness *T* as functions of *κ*_3_ (Fig. 6b and c). At *κ*_3_ = 0.1*κ*_4_, both *N*_l_ and *T* remained at values characteristic of a thin, monolayer-like sheet. As *κ*_3_ increased, *N*_l_ and *T* increased concomitantly, indicating a transition to compositionally sorted multilayers; in the data shown, this transition occurred around *κ*_3_ ≈ 0.15*κ*_4_ (Fig. 6b and c). For larger *κ*_3_, both *N*_l_ and *T* approached plateau values, indicating that further increases in *κ*_3_ produced limited additional increases in layer number and thickness within the explored range (Fig. 6b and c).

Together, these results show that the recursive assignment uses only two tension levels (*κ*_3_ and *κ*_4_), yet can be systematically extended to systems with increasing numbers of cell types to generate compositionally sorted multilayers with progressively larger numbers of layers. In principle, increasing the number of cell types under the same two-level prescription allows the design to scale to arbitrarily many layers.

## 4. Discussion

This study proposes a mechanics-based design viewpoint in which cell–cell interfacial tension serves as a tunable control handle for layered tissue architecture. While interfacial tension has long been recognized as a key determinant of morphogenetic behaviors, it has remained less clear how to translate this concept into practical rules for designing predictable layered structures. By prescribing interfacial-tension relationships using a 3D vertex model, we demonstrate that tissue architecture can be systematically shaped into monolayers, bilayers, or structurally stratified organizations, and that heterogeneous assemblies can be programmed into prescribed layered arrangements through tension-driven sorting. Importantly, the recursive scheme requires only two interfacial-tension levels (high and low); in principle, this could be sufficient to specify layered architectures with an arbitrary number of layers. Together, these findings provide a compact set of design principles that connects interface-level mechanics to tissue-scale layered architecture formation and supports rational engineering of layered tissues.

In living tissues, effective cell-cell interfacial tension reflects a balance between cortical contractility and intercellular adhesion mediated by molecules such as cadherins and the actomyosin network (Fagotto, 2014; Heer and Martin, 2017; Miroshnikova et al., 2018). Perturbations of this balance can drive diverse morphogenetic behaviors, including delamination/extrusion and cell sorting (Clarke and Martin, 2021; Devanny et al., 2021; Krieg et al., 2008). Building on this developmental mechanism, our simulations provide a complementary mechanical interpretation by treating interfacial tension as an explicit and tunable control handle for tissue architecture: in homogeneous tissues, increasing effective interfacial tension destabilizes a monolayer and drives a transition to structural stratification (Fig. 2 and Fig. 3), whereas in heterogeneous tissues, differential interfacial tensions enable out-of-plane sorting and drive the formation of segregated layers (Fig. 4, Fig. 5 and Fig. 6). Together, these results demonstrate that interfacial tension—arising from contractility-adhesion balance in vivo—can be leveraged as a controllable mechanical input to engineer predictable layered tissue organizations.

To achieve programmable control over tissue architecture, recent advances in synthetic biology and cell-surface engineering enable targeted manipulation of cell-cell interactions and their mechanical consequences (Brassard and Lutolf, 2019; Yin et al., 2016). For example, SynNotch-based circuits can implement contact-dependent regulation of adhesion molecules to steer multicellular self-organization (Almeida-Pinto et al., 2023; Gaspar et al., 2020), and coupling synthetic signaling with cadherin-mediated adhesion has been shown to generate layered assemblies (Toda et al., 2018). Complementing genetic control of adhesion, optogenetic tools provide rapid spatiotemporal control of actomyosin contractility—e.g., OptoShroom3 for light-induced apical constriction (Martínez-Ara et al., 2022) and OptoMYPT for local relaxation of contractility (Yamamoto et al., 2021)—offering a route to impose or modulate effective tension patterns. More recently, morphogen-responsive control of cadherin adhesion was shown to generate sharp 3D tissue domains (Mizuno et al., 2024). However, in such high-dimensional control schemes, it is not straightforward to identify conditions that reliably yield a desired architecture; simulation-guided design can address this gap by inferring the interfacial-tension prescriptions required for target structures, enabling a practical workflow that couples computational design with synthetic and optogenetic implementation.

While our 3D vertex simulations demonstrate that prescribed interfacial-tension relationships can generate monolayer-to-stratified-state transitions and organize multiple cell types into layered configurations, stratification in developing and engineered tissues is also shaped by additional active processes and feedbacks. For example, the orientation of cell division can contribute to basal– suprabasal boundary formation by placing daughter cells into different layers (Du et al., 2018). Active migration, delamination, and extrusion—often triggered by crowding and local remodeling of contractility—also influence vertical positioning and differentiation dynamics (Miroshnikova et al., 2018). We therefore propose that such active processes provide driving forces for vertical displacement, whereas interfacial-tension differences provide a mechanical context that biases contact rearrangements and helps stabilize layer segregation, conceptually analogous to boundary-like constraints implicated in epithelial integrity (Ohno et al., 2021). Extending the present framework to incorporate proliferation, division orientation, and additional active remodeling rules will enable more quantitative analysis of tissue morphogenesis and broaden predictive design of stratified tissues.

## 5. Conclusion

Overall, this work demonstrates the value of simulation-guided design for translating cell-scale mechanical control into tissue-scale architecture. By using simulations to map prescribed interfacial-tension relationships to emergent layered outcomes, design spaces can be explored systematically, and robust tension prescriptions can be identified before experimental implementation. Integrating such computational design with synthetic and optogenetic control of adhesion and contractility will provide a practical route to narrow experimental search spaces and to realize targeted layered architectures in engineered tissues and organoid systems.

## CRediT authorship contribution statement

**C. Thiticharoentam:** Writing – review & editing, Writing – original draft, Visualization, Validation, Investigation, Formal analysis, Data curation.

**S. Fukamachi:** Writing – review & editing, Supervision, Methodology, Investigation.

**S.A. Horiguchi:** Writing – review & editing, Writing – original draft, Supervision, Methodology, Investigation, Funding acquisition.

**S. Okuda:** Writing – review & editing, Writing – original draft, Supervision, Software, Resources, Project administration, Methodology, Investigation, Funding acquisition, Conceptualization.

## Acknowledgments

We would like to thank all the members of the Okuda laboratory for their insightful discussions and feedback. This work was supported by WISE Program for Nano-Precision, Medicine, Science, and Technology of Kanazawa University by MEXT, JST, the establishment of university fellowships towards the creation of science technology innovation, Grant Number JPMJFS2116 and JST SPRING, Japan Grant Number JPMJSP2135, Japan (to C.T.), Japan Science and Technology Agency (JST) PRESTO, Japan, Grant Number JPMJPR25KB (to S.H.), JST, CREST, Grant Number JPMJCR1921, JPMJCR24B2 (to S.O.), the Japan Society for the Promotion of Science (JSPS), KAKENSHI, Grant Number 22H05170, 24H01398, 24H01937, 25K01118, 25H01053, 25K22468 (to S.O.) and the World Premier International Research Center Initiative, MEXT, Japan (to S.O.).

## Data availability

Data will be made available on reasonable request.

